# High Performance Virtual Screening by Targeting a High-resolution RNA Dynamic Ensemble

**DOI:** 10.1101/151407

**Authors:** Laura R. Ganser, Janghyun Lee, Bharathwaj Sathyamoorthy, Aman D. Kansal, Dawn K. Merriman, Hashim M. Al-Hashimi

## Abstract

Dynamic ensembles that capture the flexibility of RNA three-dimensional (3D) structures hold great promise in advancing RNA-targeted drug discovery. Here, we experimentally screened the transactivation response element (TAR) RNA from human immunodeficiency virus type-1 (HIV-1) against ~100,000 small molecules. We used this dataset, along with 240 known hit molecules, to evaluate virtual screening (VS) against a high-resolution TAR ensemble determined by combining NMR spectroscopy and molecular dynamics (MD) simulations. Ensemble-based VS (EBVS) scores molecules with an area under the receiver operator characteristic curve (ROC AUC) of 0.87 with ~50% of all hits falling within the top 2% of scored molecules, and also correctly predicts the different TAR inter-helical structures when bound to six molecules. The prediction accuracy decreased significantly with decreasing accuracy of the target ensemble or when docking against a single RNA structure. These results demonstrate that experimentally determined ensembles can significantly enrich libraries with structure-specific RNA binders as well as motivate the continued development of methods for improving ensemble accuracy.

Accompanying the non-coding RNA (ncRNA) revolution has been growing interest in RNA as a drug target, particularly for diseases that do not have suitable protein targets^1–6^. The growing list of ncRNA targets now includes microRNAs, long ncRNAs, repetitive RNA transcripts, riboswitches, and viral RNAs. Much attention has been directed towards targeting ncRNAs with small molecules, which do not suffer from the inherent delivery limitations associated with oligonucleotide-based therapeutics, and which can be tailored to specifically target the diverse 3D structures of RNA^2–5^. There are, however, fundamental challenges in targeting RNA using small molecules^4–7^. Libraries used in high-throughput screening (HTS) are optimized for the deep, hydrophobic pockets of protein targets and not the more polar and solvent exposed pockets typical of RNA targets^7^. Screens targeting ncRNAs typically yield very few hit molecules most of which show little discrimination against other cellular RNAs, have unfavorable pharmacological properties, and poor activity in cell-based assays. Rational approaches to identify small molecules that bind specific RNA secondary structure have had some success^8^, but achieving the desired selectivity and efficacy is difficult given the reoccurrence of secondary structural motifs across the transcriptome. While many RNA targets fold into unique 3D structures, application of structure-based approaches such as computational docking to identify compounds that specifically bind to these structures has been hampered by the high flexibility of RNA and its propensity to undergo large changes upon binding to small molecules that are difficult, if not impossible, to model computationally^9^.

Recently, hybrid experimental-computational approaches have been developed that enable the determination of dynamic ensembles of biomolecules at atomic resolution^10–15^. The ensembles describe the different conformations sampled by a given biomolecule in solution. Studies on both proteins and nucleic acids have shown that conformations similar to those observed in ligand bound states are often significantly populated within the ensemble of unbound conformations^13–16^. Inspired by these discoveries, we^9^ and others^17^ recently proposed to carry out EBVS by subjecting all conformers in experimentally determined ensembles to computational docking^9,17^. Our approach was demonstrated using a dynamic ensemble of HIV-1 TAR^18^ determined using two sets of NMR residual dipolar coupling (RDC) data and MD simulations^9,13^. RDCs provide long-range information regarding the orientation of individual bond vectors in biomolecules relative to an alignment frame and are sensitive to internal motions spanning picosecond-to-millisecond timescales^19,20^. EBVS was able to score a small set of 38 known TAR binders with an accuracy comparable to that obtained when docking 48 small molecules to their known bound RNA structure^9^. Experimentally testing the top 57 scoring small molecules out of a screen of 51,000 compounds yielded six hit molecules that bind TAR *in vitro*, including the first example of a small molecule that binds an RNA apical loop, and a highly selective compound that inhibits HIV replication in an HIV indicator cell line with IC50~20 μM^9^.

EBVS can in principle provide a basis for pre-screening compound libraries to enrich a subset of molecules with potentially active hits that specifically bind to the 3D structure(s) of an RNA target. EBVS could also aid exploration of new regions of chemical space unhindered by physical constraints posed by existing libraries to identify novel RNA-binding scaffolds. To rigorously test the utility of EBVS in such pre-screening applications, we first generated a large dataset by subjecting HIV-1 TAR (Fig. 1a) to experimental HTS against ~100,000 drug-like organic molecules (Methods and Supplementary Fig. 1-4). This represents one of the largest RNA-small molecule screens reported to date. The screen employed a Tat peptide displacement assay (Z score=0.71), included several steps to prevent false positives and false negatives (see Methods), and yielded a total of seven new hits (Supplementary Table 1) representing two novel small molecule classes that bind TAR and inhibit the TAR-Tat interaction (Supplemental Fig. 1). Such low hit rates (~0.007%) are typical for large RNA-small molecule screens^7^.

**Figure 1.**
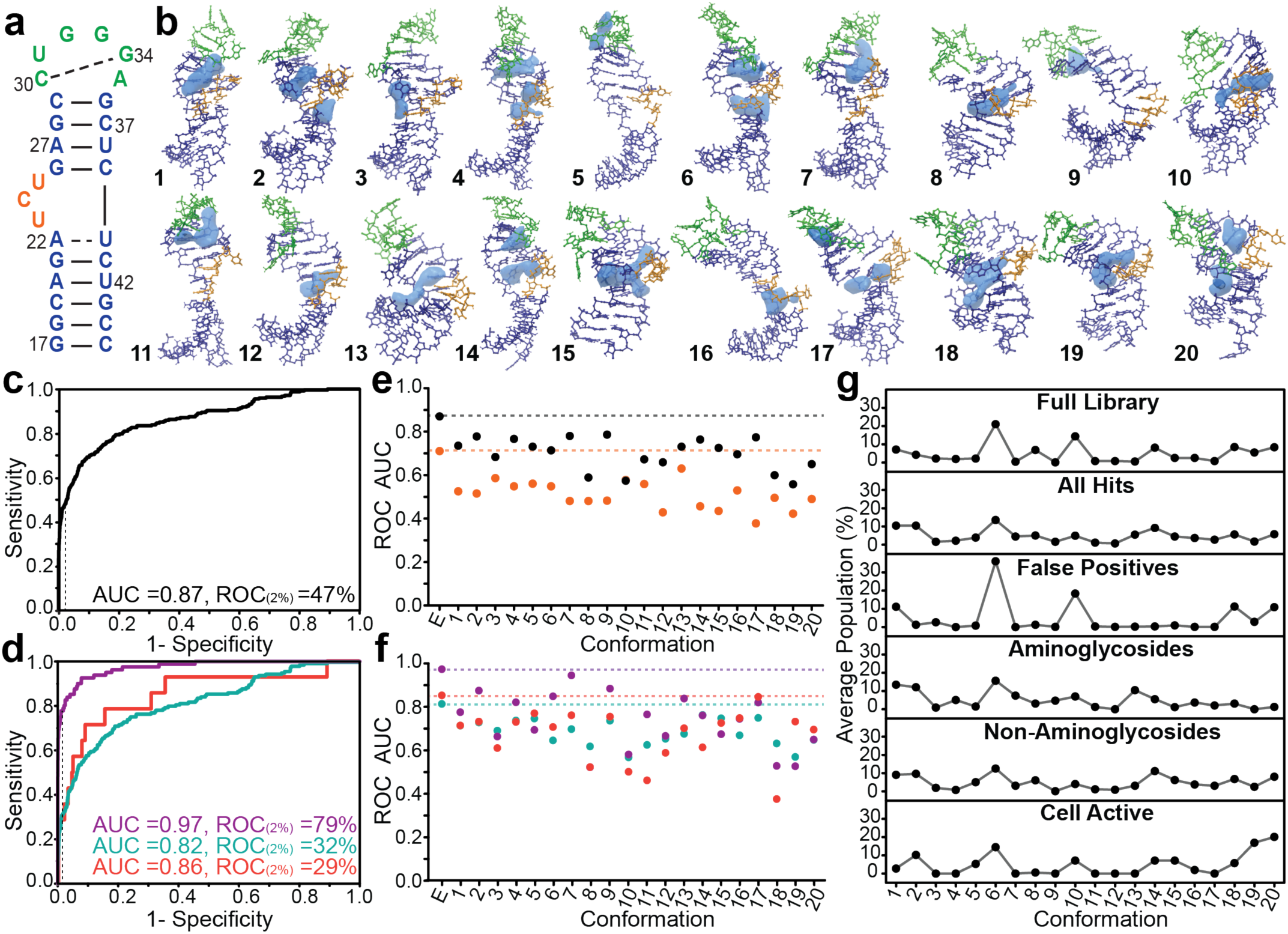
Evaluating EBVS against HIV-1 TAR using an experimental dataset derived from HTS using ~100,000 small molecules. **a.** Secondary structure of HIV-1 TAR. **b.** Ensemble of HIV-1 TAR (E0) determined using four sets of RDCs showing binding pockets (blue) predicted by ICM. **c-d.** ROC curve analysis of EBVS showing enrichment of the **c.** 247 hits and **d.** aminoglycoside hits (purple), non-aminoglycoside hits (blue), and cell-active hits (red). Dotted lines indicate the ROC_(2%)_ values. **e-f.** EBVS ROC AUC values compared to ROC AUC values obtained when docking to the individual conformers of **e.** the parent ensemble E0 (black) or random ensemble E3 (orange) and **f.** for specific hit molecule categories of the parent ensemble: aminoglycosides (purple), non-aminoglycosides (blue), and cell-active compounds (red). **g.** Docking predicted Boltzmann populations for each conformer when bound to a small molecule averaged over all molecules in the library. Values shown for all 247 hit molecules, false positive molecules, aminoglycoside hits, non-aminoglycoside hits, and cell-active hits.

The ~100,000 molecule library used in HTS was augmented with an additional 240 molecules previously shown to bind HIV-1 TAR (Supplementary Table 2) and the combined library used in EBVS. The additional hits included 81 aminoglycosides and aminoglycoside conjugates as well as a diverse set of 159 small molecules including derivatives of beta-carboline, quinolone, diphenylfuran, phenothiazine and many others. This is by far the largest and most diverse set of experimentally verified binders and non-binders used to evaluate VS against any RNA target.

EBVS was carried out by computationally docking the virtual small molecule library to a recently reported high-resolution dynamic ensemble of HIV-1 TAR RNA^14^. The ensemble contains twenty conformers that describe the various TAR conformations sampled in the unbound state (Fig. 1b)^14^. Compared to the previous TAR ensemble used in VS^9,13^ (see E1 below), this improved ensemble was determined using four rather than two sets of RDCs, obtained using a variable-elongation approach^21^. In addition, a longer MD simulation (8.2 μs versus 80 ns) was used to generate the pool of starting TAR conformations^14^. Using the docking program Internal Coordinate Mechanics (ICM)^22^, each small molecule was docked against each of the twenty TAR conformers using binding pockets defined by the ICM PocketFinder module (Fig. 1b). The docking scores computed for a given small molecule across the twenty TAR conformers were averaged assuming a Boltzmann distribution (see Methods).

To evaluate the ability of EBVS to enrich true hits while discriminating against non-hits, we used the EBVS ranked scores to compute a ROC curve, which compares the fraction of hits correctly identified (sensitivity) versus the fraction of non-hits incorrectly selected (1-specificity) at all possible docking score cutoffs. The ROC AUC describes the global enrichment of true binders with AUC=1.0 representing perfect enrichment and AUC=0.5 representing random selection of hits and non-hits. Strikingly, EBVS globally enriches the compound library (247 hits and ~100,000 non-hits) with a ROC AUC=0.87 (Fig. 1c). We find strong early enrichment such that we would have identified 47% of hits after screening only 2% of non-hits (ROC_(2%)_=47%) (Fig. 1c). This corresponds to a hit rate of 5.6% compared to 0.2% had the whole library been screened. This level of enrichment was robustly observed across the different categories of experimentally verified hits (Fig. 1d) including aminoglycosides, non-aminoglycosides, and compounds with demonstrated cell-based activity (Supplementary Table 2). These levels of enrichment are comparable to best-case results when docking to known bound structures of RNA or protein^23,24^.

By comparison, docking to individual TAR conformers either from the NMR ensemble (AUC= 0.56-0.79) or selected randomly (see E3 below) from the MD generated pool (AUC= 0.38-0.63) leads to a precipitous drop in enrichment (Fig. 1e). This trend is observed across all molecule categories (Fig. 1f) and highlights the importance of docking to an experimentally informed ensemble. Docking predicts relatively uniform binding to the twenty TAR conformers, indicating that all conformers in the ensemble contribute to the docking scores across the library. Interestingly, we observe conformer-specific enrichment in subsets of molecules within the library (Fig. 1g). For example, conformers 1 and 2 enrich for both aminoglycoside and non-aminoglycoside hits, conformer 13 enriches for aminoglycoside hits, and conformers 19 and 20 enrich for the 14 cell-active hits.

Next, we asked whether the accuracy of the TAR ensemble was an important determinant of the docking predictions. We repeated EBVS using four additional (E1-E4) twenty-member TAR ensembles that have variable levels of accuracy as evaluated based on their ability to predict the four NMR RDC datasets^14^ (Fig. 2a). The parent ensemble (E0) predicts the four RDC datasets with RMSD = 4.0 Hz^14^. By comparison, E1 was determined using only two RDC datasets and a shorter MD simulation and predicts all four RDC datasets with RMSD = 7.1 Hz^13^. E2 and E3 were constructed by randomly selecting conformers from two different MD pools (RMSD = 8.6 and 11.0 Hz, respectively) while E4 was deliberately selected (see Methods) to poorly satisfy the RDCs (RMSD = 18.0 Hz). Significantly, there was a strong correlation between the quality of the ensemble and the EBVS enrichment as measured by the ROC AUC (Fig. 2b) and ROC(2%) (Fig. 2c) values. These trends were generally observed across different types of hits (Fig. 2d-e).

**Figure 2.**
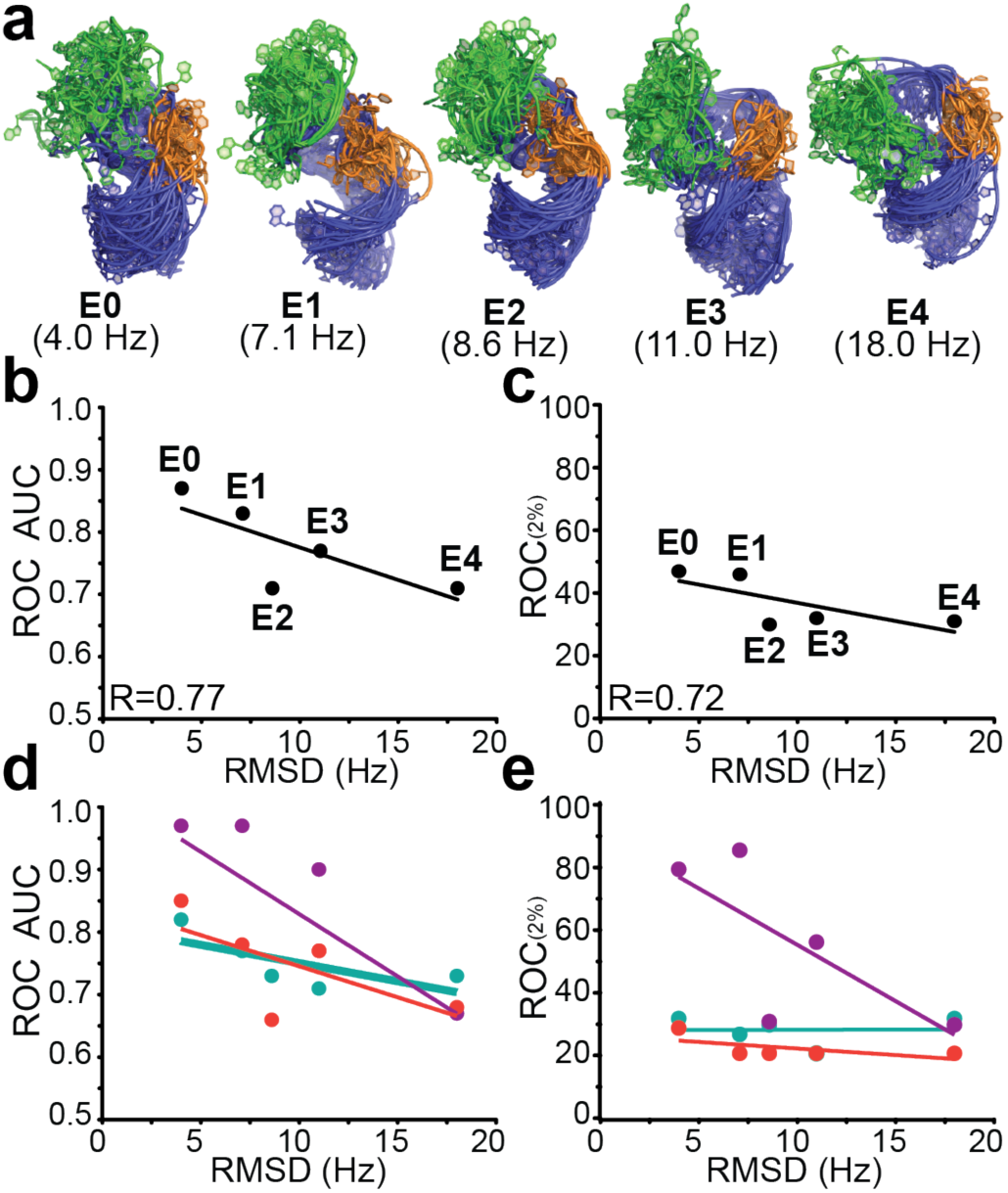
The accuracy of EBVS predictions is directly correlated to the quality of the HIV-1 TAR ensemble. **a.** Distinct ensembles of unbound TAR with variable accuracy as assessed based on agreement with experimental RDCs (RMSD shown in parentheses). **b-e.** Correlation between the TAR ensemble accuracy (RDC RMSD) and **b.** ROC AUC values for all 247 hits; **c.** ROC AUC values for aminoglycoside hits (purple), non-aminoglycoside hits (blue) and cell-active hits (red); **d.** ROC_(2%)_ values for all 247 hits; and **e.** ROC_(2%)_ values for aminoglycoside hits (purple), non-aminoglycoside hits (blue) and cell-active hits (red).

Next, we asked whether EBVS correctly predicts the small molecule bound TAR conformations for six TAR complexes for which high-resolution NMR structures have previously been reported: arginine (1ARJ), acetylpromazine (1LVJ), neomycin B (1QD3), RBT203 (1UUD), RBT158 (1UUI), and RBT550 (1UTS). Figure 3a compares the NMR structures to the EBVS top-scoring conformations over twenty repeated docking runs. While comparison of structures is complicated by the fact that TAR is known to retain a high degree of conformational flexibility when bound to small molecules^25–27^, qualitatively, we observe good agreement between the NMR and the EBVS-predicted bound TAR conformations (Fig. 3a). We find more variability in the ligand placement for docking than for NMR structures, but in general, docking correctly places molecules near the bulge-expanded major groove (Fig. 3a). The one exception is neomycin B; docking predicts binding to either the major or minor groove of the bulge or upper stem whereas it binds to the minor groove of the lower stem in the NMR structure, which is unusual since most aminoglycosides bind the RNA major groove^28^.

**Figure 3.**
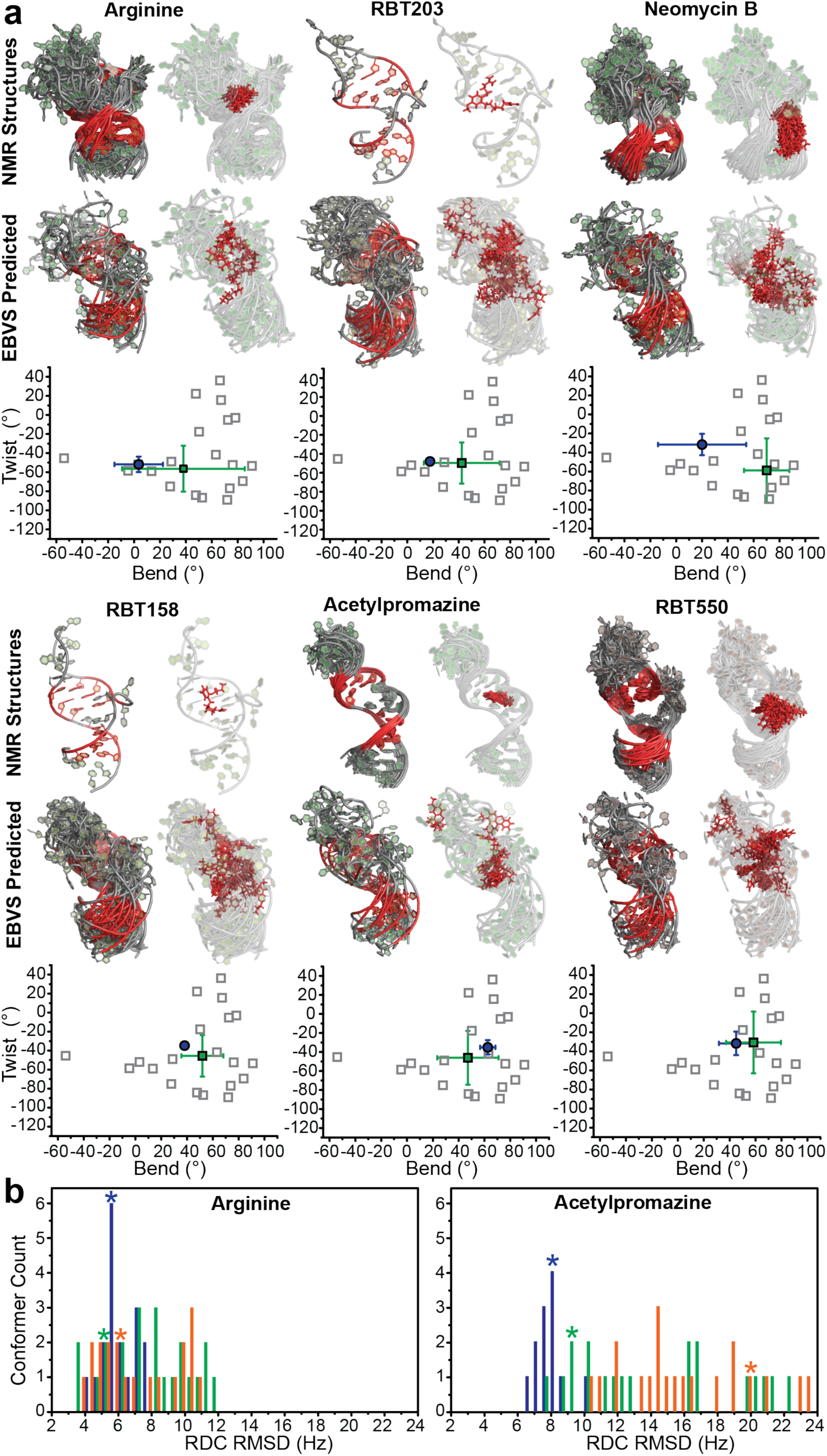
Assessing EBVS-predicted small molecule bound TAR conformations. **a.** For each small molecule, NMR structures (all models) are compared to EBVS-predicted structures (all conformations predicted to be > 25% populated following 20 independent docking runs). Colored in red are the base pairs in the upper and lower stems used to superimpose the RNA structures (left) and the small molecule (right). Also shown are the inter-helical bend (β_h_) and twist (α_h_+γ_h_) angles^30^ (negative and positive twist angles correspond to over- and under-twisting respectively) averaged over all models in the NMR structure (blue circles) and the Boltzmann-weighted angles predicted by EBVS averaged over 20 docking runs (green squares). Also shown are the angles for individual conformers in the parent ensemble (open squares). **b.** Histograms of the calculated RDC RMSD (Hz) between ligand-bound TAR RDCs and the NMR structures (blue), E0 conformers (green), and E3 conformers (orange). The asterisks indicate the RDC RMSD for the lowest energy NMR structure (blue) and the EBVS-predicted docking structure for E0 (green) and E3 (orange) for docking with thoroughness set to 20 over 5 docking runs.

Studies have shown that ncRNAs can adopt structures that vary with respect to the global orientation of helical domains when bound to different small molecules^18,25,29^. We therefore examined if EBVS correctly predicts the variations in the TAR inter-helical structure when bound to the six small molecules. We compared the inter-helical Euler angles (α_h_ β_h_ γ_h_)^30^ describing the bend (β_h_) and twist (α_h_+γ_h_) angles between the two TAR helices. For all molecules, the EBVS predicted inter-helical Euler angles are in quantitative agreement with those observed in the bound NMR structures (Fig. 3a). The EBVS predicted TAR structures are biased towards the NMR bound structures relative to all twenty conformers in the parent E0 ensemble, indicating that docking enriches for accurate bound TAR conformations (Fig 3a). In contrast, the predicted bound structures obtained by applying EBVS to the randomly selected E3 ensemble show significantly reduced agreement with the NMR structures particularly for RBT550, RBT158 and acetylpromazine (Supplementary Fig. 6). Similar results were obtained when increasing the thoroughness used in docking (Supplementary Fig. 7).

We confirmed these findings by evaluating the agreement between the EBVS-predicted structures and RDCs previously measured for two of the TAR-small molecule complexes (argininamide^26^ and acetylpromazine^27^). These RDCs were not used in the NMR structure determination. This analysis could not be performed for neomycin B, for which there are also available RDCs, because docking predicted more than one small molecule bound conformation, which complicates analysis of the RDCs (see Methods). The docking predicted structures satisfy the RDCs to a comparable degree as do the high-resolution NMR structures of the TAR complexes (Fig. 3b and Supplementary Table 3). Applying EBVS to the randomly selected E3 ensemble shows less favorable agreement with the measured RDCs of the bound complex, particularly for acetylpromazine (Fig. 3b). These results indicate that EBVS can allow the identification of small molecules that target specific RNA conformations within an ensemble, potentially enabling targeted conformational modulation of RNA with small molecules^5^.

In conclusion, EBVS significantly enriches true TAR hits including compounds with documented cell-based activities out of a large pool of ~100,000 non-hits. This result is unlikely to be coincidental since we observe that docking predicts bound conformations that agree with experimental data including NMR structures and NMR RDCs of TAR-ligand complexes. Finally, we show a direct correlation between the accuracy of the unbound TAR ensemble and the quality of docking predictions. This suggests that structure-based targeting of flexible biomolecules is possible with accurate ensemble representations, motivating the continued development of experimentally determined dynamic ensembles and methods of evaluating their accuracies. While we have demonstrated a specific application of EBVS to HIV-1 TAR, further studies are needed to evaluate how predictions might vary when using different docking programs, starting structures in MD simulation, MD force field, and complexity of the target RNA. Nevertheless, EBVS holds great promise for enriching compound libraries used in RNA-small molecule screening and can thus be utilized to maximize success in RNA-targeted drug discovery.

## AUTHOR CONTRIBUTIONS

Experiments were designed by L.R.G, J.L. and H.M.A.-H., performed by L.R.G. J.L., B.S. A.K. and D.K.M, and analyzed by L.R.G., L.R.G. and H.M.A.-H. wrote the manuscript.

## ACKNOWLEDGEMENTS

We would like to thank the University of Michigan Center of Chemical Genomics, in particular Martha J. Larsen and Steve Vander Roest, for their help in managing the small molecule library and carrying out high throughput screening. We would also like to thank the Duke Magnetic Resonance Spectroscopy Center for their resources and assistance in carrying out NMR measurements. Finally, we acknowledge the Duke Compute Cluster for computational resources and support. This work was supported by the US National Institutes of Health [P50 GM103297, R01 AI066975 to H.M.A.-H. and T32 GM08487 to L.R.G].

## ONLINE METHODS

### Library composition

The small molecule library used in experimental HTS consisted of 103,498 drug-like small molecules that were available at the Center for Chemical Genomics (CCG), University of Michigan, Ann Arbor. 100,000 molecules were synthetic organic molecules with drug-like properties purchased from ChemDiv (http://www.chemdiv.com). The other 3,498 compounds consisted of 2,000 bioactive molecules from MicroSource Discovery Systems Inc. (http://www.msdiscovery.com), 446 molecules from the National Institute of Health clinical collection, and 1052 molecules that the CCG had previously found to be active against other targets. The library was stored as stock solutions of 2-5 mM molecule in DMSO for ~3 years for initial screens. Repurchased molecules were stored as stock solutions of 3-20 mM compound in DMSO for ~1 year with the exception of CCG-39701 which was stored as a powder and dissolved in water before use.

A virtual library of the same 103,498 molecules was downloaded from the CCG and saved in sdf file format. The library was enriched with 240 molecules drawn using ChemDraw (CambridgeSoft) and saved in sdf file format that were previously reported to bind TAR *in vitro* (Supplementary Table 2). These molecules were identified through a literature search of TAR binders. Peptide and metal-binding molecules were excluded. For papers reporting many chemically similar molecules, only the tightest binding derivatives were included to maintain chemical diversity in our library. The protonation states of small molecules were predicted at pH 7.4 using the Major Microspecies module in the Calculator Plugin (ChemAxon).

### Preparation of HIV-1 TAR RNA

HIV-1 TAR RNA (Fig. 1a) for NMR studies and experimental assays was prepared by *in vitro* transcription using DNA template containing the T7 promoter (Integrated DNA Technologies). DNA template was annealed at 50 μM DNA in 3 mM MgCl2 by heating to 95°C for 5 min and cooling on ice for 30 min. The transcription reaction was carried out at 37°C for 12 hours with T7 RNA polymerase (New England BioLabs) in the presence of ^13^C^/15^N labeled or unlabeled nucleotide triphosphates (Cambridge Isotope Laboratories, Inc). RNA was purified using 20% (w/v) denaturing polyacrylamide gel electrophoresis (PAGE) with 8 M urea and 1X TBE. Purified RNA was extracted from the gel by electroelution in 1X TAE buffer and purified by ethanol precipitation. Purified RNA was dissolved in water to 50 μM RNA, heated to 95°C for 5 min and cooled on ice for 1 hour to anneal. For NMR experiments, ^13^C^/15^N labeled RNA was exchanged into phosphate NMR buffer [15 mM NaH2PO4/Na2HPO4, 25 mM NaCl, 0.1 mM EDTA, 10% (v/v) D2O at pH 6.4]. For *in vitro* assays, unlabeled RNA was diluted to 150 nM in Tris-HCl assay buffer [50 mM Tris-HCl, 50 mM KCl, 0.01% (v/v) Triton X-100 at pH 7.4].

### Preparation of HIV-1 Tat peptide

The Tat peptide used in HTS, (5-FAM)-AAARKKRRQRRRAAA-Lys(TAMRA), was purchased from LifeTein (Hillsborogh, NJ). The peptide was purified by HPLC and analyzed by Electrospray Ionization Mass Spectrometry; purity was > 95%. The peptide was stored at -20°C as a 100 μM stock solution in Tris-HCl assay buffer and diluted to 60 nM with Tris-HCl assay buffer for use in HTS.

### High throughput screening assay

HTS utilized a previously described TAR-Tat displacement assay^31^ that relies on the fact that the Tat peptide is highly flexible when free is solution and becomes structured upon binding to TAR^32–34^. When the Tat peptide is flexible, its two terminal fluorophores, fluorescein and TAMRA, interact and their fluorescence is quenched. Alternatively, in its extended form bound to TAR, the fluorophores are held at a distance allowing fluorescence resonance energy transfer (FRET) from fluorescein to TAMRA. Thus, as inhibitor displaces Tat, there is a decrease in fluorescence signal (excitation: 485 nm, emission: 590 nm).

### High throughput screening

The workflow for HTS is shown in Supplementary Figure 1. The full library was tested in a primary screen using a single point measurement (N=1) and 260-fold excess molecule [50 nM TAR, 20 nM Tat, and 13 μM molecule] followed by a confirmation screen of triplicate measurements (N=3) for the 2812 molecules that showed activity, defined as a change in fluorescence signal three standard deviations above the negative control (Tat alone). Molecules were pin-tooled (200 nL) into opaque 384-well microplates by Biomek FX 384-well nanoliter HDR (Beckman) and Mosquito X1 (TTP Labtech). TAR and Tat were dispensed with Multidrop reagent dispenser (Thermo Scientific). Assay mixtures were incubated at room temperature for 10–15 minutes prior to fluorescence measurements using a Pherastar plate reader (BMG Labtech). Each plate during HTS contained 16 wells of TAR and Tat without molecule to serve as the negative control and 16 wells of Tat only to serve as the positive control. With these controls, the Z-factor^35^ was calculated for each microplate; the average Z-factor throughout the screening campaign was 0.71.

### Dose Response Assays

A total of 267 molecules with reproducible activity were tested in a dose response assay and those with IC50 values below 100 μM were considered hits. Dose-response assays were performed such that the final assay concentrations were 50 nM TAR, 20 nM Tat, and 1-1000 μM molecule in Tris-HCl assay buffer. Assays were performed in parallel with and without 100-fold excess bulk yeast tRNA to test specificity and in the absence of RNA (Tat only) to measure background signal. There were 137 molecules that caused fluorescence intensity change with Tat alone, suggesting they bound Tat; these were removed from further analysis. Assays were performed in opaque 384-well microplates and read with a Clariostar plate reader (BMG Labtech). Fluorescence signal was normalized to the highest intensity after subtracting background signal. Dose response curves were fit to Equation 1 with OriginPro (OrginLab) using the instrumental weighting method. Equation 2 was used to obtain IC50 values.

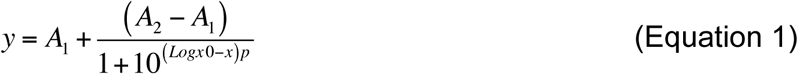

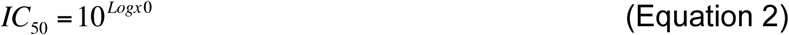

where A_1_ and A_2_ are the lowest and highest signals, respectively; p is the hill slope; and log(x_0_) is the logarithm to base 10 of the concentration at half response. All variables were allowed to float during the fit. Assays were measured in triplicate and the standard deviation is reported.

### Validation of Hits

The 17 small molecule hits from the dose response assays were re-purchased and re-tested for activity in addition to 56 molecules with chemical similarity to these hits, defined as having >80% similarity based on sphere exclusion clustering performed with JKlustor package (ChemAxon). Next, 32 molecules, including all 17 hits and 15 chemically similar molecules with possible activity in the assay, were tested for TAR binding by NMR chemical shift titrations employing [^13^C-^1^H] SOFASTHMQC NMR experiments^36^. NMR experiments were performed at 298 K on 600 MHz and 800 MHz Agilent spectrometers equipped with triple-resonance HCN cryogenic probes.^13^C/^15^N-labeled TAR was exchanged into NMR buffer. Concentrated stocks of molecule in DMSO were added to TAR such that no more than 10% (v/v) DMSO was added to the buffer. Free TAR controls had equivalent volumes of DMSO to compensate for minor changes that may be induced by DMSO. Spectra were processed using nmrPipe^37^ and SPARKY^38^.

Nine molecules were inactive in both the displacement assay and NMR when retested with fresh molecule, suggesting that the original activity was due to contamination or degradation. One of the 56 molecules with chemical similarity to the hits was active in both the displacement assay and NMR, despite not being identified as a hit in the primary screen. Three molecules had activity in the assay, but did not bind based on NMR chemical shift titrations. Inspection of the Tat-only control for these molecules suggest that they likely bind Tat rather than TAR in the displacement assay (Supplementary Fig. 5). These should have been identified earlier in the workflow, but the fluorescence change in the presence of Tat may not have been large enough. Overall, seven molecules were confirmed to bind TAR RNA based on their activity in the TAR-Tat displacement assay (Supplementary Fig. 1b) and their ability to induce chemical shift perturbations in the TAR NMR spectra (Supplementary Fig. 1c).

### HTS hit molecules

The seven novel confirmed HTS hits fall into two general scaffolds; five of the hits share an anthraquinone scaffold with two sites of derivatization while the other two molecules share similar geometries with napthyl and quinazoline cores (Supplementary Table 1). All anthraquinone derivatives maintain their IC50 in the presence of 100-fold excess tRNA, suggesting they are specific for TAR (Supplementary Table 1). Several additional anthraquinone molecules with minor differences to the five hits did not bind TAR (Supplementary Fig. 2). The fact that small structural changes can ablate binding is consistent with the hit molecules making specific interactions with TAR. Interestingly, the anthraquinone hits and chemically similar molecules exhibited a change in color from orange to blue when diluted from 100% DMSO to an aqueous solution, likely due to DMSO reacting with the anthraquinone to form DMSO-anthraquinone, as described previously^39^. All experiments were performed with the derivatives in the blue state. The second class of molecules has much poorer activity in the presence of tRNA, suggesting that they are not specific for TAR (Supplementary Table 1). This lack of specificity is likely due to the positive charges on these molecules, which would interact non-specifically with the RNA backbone.

The addition of the small molecule hits to TAR resulted in large chemical shift perturbations or line broadening in several residues throughout TAR (Supplementary Fig. 1c-f). As expected, hits with similar chemical structures induce similar chemical shift perturbations indicating that they interact with TAR using similar binding modes (Supplementary Fig. 1c-d). There are however two interesting exceptions. One of the five anthraquinone molecules, CCG-133905, induces significantly more broadening consistent with tighter binding, although other factors such as aggregation could induce the broadening (Supplementary Fig. e). The anthraquinone molecule, CCG-133994, which contains an ester and an amine, induces chemical shift perturbations that are distinct from the other anthraquinone molecules, suggesting a distinct binding mode for this molecule (Supplementary Fig. f). Furthermore, NMR reveals that this molecule is in slow exchange, which is in agreement with the fact that it is the tightest binder in the TAR-Tat displacement assays.

### Identification of false negatives in HTS

Molecules that were insoluble in the assay conditions or that had fluorescence interference with the Tat peptide would have been excluded from the HTS workflow and this may represent a possible source of false negatives in our dataset. To investigate this possibility, we selected nine molecules in the top ~3% of docking scores and tested them for TAR binding using NMR. Three of the nine molecules did in fact bind TAR under NMR conditions (Supplementary Fig. 4b). Closer analysis revealed different factors led to the exclusion of these small molecules from HTS during the primary screen. One molecule, CCG-39701, was insoluble in DMSO but was active in the assay when dissolved in water (Supplementary Fig. 4a). The other two molecules, CCG-208298 and CCG-100975, had fluorescence interference at high concentration preventing determination of an accurate IC50 (Supplementary Fig. 4a). These results highlight a weakness of HTS, which is the presence false negatives, which we recognize is a source of uncertainty in our analysis. To avoid biasing the results, these molecules were not included in the ROC analysis.

### Virtual Screening

VS was performed using the docking program Internal Coordinate Mechanics (ICM, Molsoft)^22^. Docking was set up as described previously^19^. Briefly, each of the 20 conformers of the TAR dynamic ensemble^34^ was uploaded to ICM in PDB format and converted to ICM objects using the default options (waters deleted and hydrogens optimized). Binding pockets were identified with the ICM PocketFinder Module using a tolerance value of 4.6. Receptor maps were generated to include all atoms within 5 Å of the predicted binding pockets with atom occupancy weighted. Docking was run with a thoroughness value of 1, flexible ring sampling level 2, and covalent geometry relaxed. For high-thoroughness docking, the thoroughness value was increased to 20, and all other parameters remained the same.

### Ensemble-Based Docking Scores

The docking scores provided by ICM represent predicted binding energies in kcal/mol. For each molecule, the fractional population of all 20 TAR conformers was calculated using the Boltzmann distribution (Eq. 3). The population of each conformer was multiplied by its docking score and these values were summed over all conformers to calculate the population-weighted score of each molecule (Eq. 4).

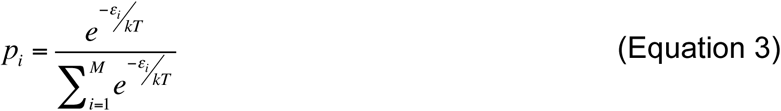

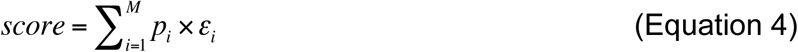

Where p_i_ is the population of conformer i, ε_i_ is the docking score of conformer i, k is the Boltzmann i constant, T is temperature (298 K), and M is the number of conformers in the ensemble.

### Receiver Operator Characteristic Curves

OriginPro (OriginLab) was used to plot receiver operator characteristic curves using Equations 5 and 6 and to calculate the area under the curve using the trapezoidal rule. Details on these equations can be found at (http://www.originlab.com/doc/Origin-Help/ROCCurve-Algorithm).

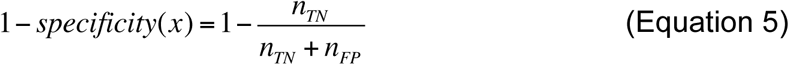

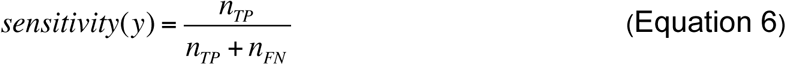

where n is the number of true negatives (TN), true positives (TP), false negatives (FN) or false positives (FP) at any given score threshold value.

### TAR Ensembles of Variable Accuracy

The RDC-derived TAR ensembles (E0 and E1) were determined as reported previously^13,14^. The randomly selected ensembles (E2 and E3) were constructed by using a random number generator to randomly select 20 structures from the two pools of TAR conformations generated using MD simulations^13,14^ containing 10,000 and 80,000 conformations, respectively. An ensemble that poorly agrees with all four RDC data sets (E4) was generated using a sample and select (SAS) Monte Carlo selection scheme to maximize the χ^2^ function assessing the agreement between measured and predicted RDCs (Eq. 7)^13^.

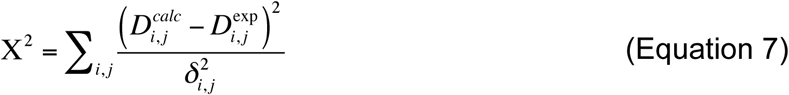

Where *i* runs over all the RDCs measured for the different constructs *j* and δ is the weight used to normalize different RDC data sets, and is set at one tenth of the range of RDCs measured for each TAR construct14. D^exp^ are the experimentally measured RDCs and D^calc^ are the predicted RDCs that were calculate by PALES^40,41^ as described below.

The quality of the various TAR ensembles used in this study was determined by evaluating how well they agree with four sets of RDC data measured on variably elongated TAR RNA molecules as described previously^14^. Briefly, the program PALES^40,41^ was used to calculate predicted RDCs based on the structures in the ensemble, after *in silico* elongation as described previously^14^. A scaling factor was used to account for variations in experimental conditions. The predicted RDCs are averaged for all structures of the ensemble assuming equal probabilities (Eq. 8).

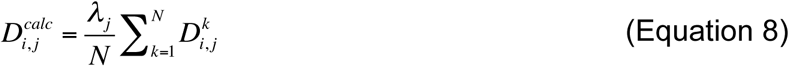

where *k* runs over the *N* conformers of the ensemble, λ_j_ is the scaling factor for the j^th^ TAR construct and D_i,j_ is the i^th^ coupling in the j^th^ construct. These calculated RDCs were then compared to measured RDCs and the RMSD (Hz) was calculated.

### Docking Predicted Pose Analysis

Pose analysis was carried out for six TAR binders for which there are NMR structures deposited in the PDB: arginine (1ARJ), acetylpromazine (1LVJ), neomycin B (1QD3), RBT203 (1 UUD), RBT158 (1UUI), and RBT550 (1UTS)^28,42–45^. For each of these molecules, docking was repeated twenty times with thoroughness set to one (Fig. 3a). It was also repeated five times with the thoroughness value increased to twenty, meaning the time spent docking each ligand was increased twenty-fold (Supplemental Fig. 7). The inter-helical angles (α_h_, β_h_, γ_h_) were computed for each ensemble conformer as well as for each model of the NMR structures using an in-house software as described previously^29^. For this calculation, the lower helix was defined as C19-G43, A20-U42 and G21-C41 and the upper helix was defined by G26-C39, A27-U38 and G28-C37. For docking results, the inter-helical angles were population-weighted based on docking scores and averaged over all runs. The inter-helical angles of the NMR structures were averaged over all models. Repeating this analysis using five docking runs with increased thoroughness improved the agreement between inter-helical angles for RBT158, RBT550, acetylpromazine and arginine, had little effect in the case of neomycin B, and worsened the agreement for the twist angle in the case of RBT203 (Supplementary Fig. 7). Interestingly, neomycin B and RBT203 were the least consistent in conformer selection across the 5 runs of docking.

We also compared the docking predicted poses and NMR structures to published RDCs of the ligand-TAR complex for arginine^26^ and acetylpromazine^27^. This analysis was done only for the most favored conformer from high-thoroughness docking over the 5 runs: conformer 8 for arginine (>86% populated in 4/5 runs) and conformer 3 for acetylpromazine (>72% populated in 4/5 runs). For the random ensemble, E3, conformer 4 was selected for arginine (>99% populated for 4/5 runs) and conformer 3 was selected for acetylpromazine (>99% populated for 5/5 runs). Likewise, only the lowest energy NMR structure was used for this analysis. The program RAMAH^46^ was used to compare these structures to the published RDC data sets. Briefly, RAMAH determines the best-fit order tensor for a given structure and set of RDCs using single value decomposition, it then uses this order tensor to calculate RDCs and reports the RMSD (Hz) between calculated and experimental RDCs. RDCs of N-H bond vectors and for residues in the bulge, loop, or terminal ends were excluded.

